# Arp2/3 complex contributes to the actin-dependent uptake of *Aspergillus terreus* conidia by alveolar epithelial cells

**DOI:** 10.1101/2025.05.21.655299

**Authors:** Natalia Mach, Julien Polleux, Lea Heinrich, Lukas Lechner, Iryna Levytska, Cornelia Lass-Flörl, Susanne Perkhofer

## Abstract

*Aspergillus terreus* is an opportunistic fungal pathogen associated with high mortality rates and intrinsic resistance to amphotericin B. Its ability to persist within host tissues without inducing strong immune responses was suggested to contribute to poor clinical outcomes. The cellular mechanisms underlying *A. terreus* interactions with host cells remain largely unexplored.

In this study, we have used a micropattern-based infection model to investigate the early interactions between *A. terreus* conidia and alveolar epithelial cells, focusing on the role of Arp2/3-dependent actin remodeling. This system allows quantitative analysis of conidia-cell interactions under defined spatial conditions. We show that *A. terreus* conidia rapidly bind to micropatterned A549 cell islands, with conidial numbers increasing over time. Conidia were found in actin- and Lamp1-positive vesicles already after one hour of infection. Inhibition of the Arp2/3 complex significantly impaired conidial binding and disrupted the formation of actin-positive vesicles, confirming the essential role of Arp2/3-mediated actin remodeling in early stages of conidial uptake. A subset of internalized conidia was localized to Lamp1-positive phagolysosomes and accumulated over time. Interestingly, we have identified a small but consistent population of phagolysosomes decorated with actin patches, potentially resembling actin flashes. These structures were entirely abolished upon Arp2/3 inhibition, indicating active cytoskeletal remodeling at the phagolysosomal interface.

Our findings provide the first mechanistic insights into *A. terreus* internalization by alveolar epithelial cells and establish Arp2/3-mediated actin dynamics as a key process in early host-pathogen interactions. This cellular pathway may contribute to intracellular persistence and help understand the delayed onset of *A. terreus* infections.

## Introduction

Fungal diseases represent a significant global health burden, particularly due to emerging antifungal resistance and high mortality rates among immunocompromised populations [1]. Currently, over 600 fungal species cause human infections, with Aspergillus spp. alone being responsible for approximately 70% of fungal-associated deaths [2]. *Aspergillus terreus* is an opportunistic fungal pathogen that causes invasive infections with high mortality, particularly in immunocompromised patients [3,4]. Although less frequently isolated than *Aspergillus fumigatus, A. terreus* is responsible for up to 15% of invasive bronchopulmonary aspergillosis (IBPA) cases and is associated with poor clinical outcomes [5–8]. A major challenge in treatment is the intrinsic resistance of *A. terreus* to amphotericin B, which limits therapeutic options and complicates prophylaxis in high-risk patients [9–12].

Upon inhalation, *Aspergillus* conidia are typically recognized and internalized by phagocytic cells through a well-coordinated mechanism involving Dectin-1, mannose receptors, and Toll-like receptors, which recognize pathogen-associated molecular patterns on the conidia surface [13–15]. Once bound, these interactions trigger the engulfment of the conidia into phagosomes, which then fuse with lysosomes to form phagolysosomes where the pathogens are supposed to be degraded [16–18]. The internalization and subsequent acidification of phagolysosomes are critical for the destruction of most fungal pathogens. However, *Aspergillus* species have developed mechanisms to resist or evade immune-mediated destruction, including melanin-mediated masking of immunogenic epitopes and inhibition of phagolysosome acidification [19,20]. Previous studies demonstrated that *A. terreus* conidia has a distinct immune evasion strategy compared to *A. fumigatus*, characterized by prolonged dormancy within the phagolysosomes of macrophages [16,21,6].

*A. terreus* conidia were shown to be rapidly phagocytosed, but remain dormant and viable within mature, acidified phagolysosomes without germination, thus avoiding immune activation [16,21]. In contrast to *A. fumigatus, A. terreus* conidia did not inhibit phagolysosome acidification and were not inactivated by low pH [19,16]. Together, these features of *A. terreus* are characterized as a dormancy-based “sit and wait” strategy likely contributing to chronic infection and poor clinical outcomes [16,6].

Previous studies showed that lung epithelial cells, even being classically non-phagocytic cells, internalize *A. fumigatus* conidia using the F-actin- and Arp2/3-dependent phagolysosomal pathway [22–26]. A study by Slesiona et al found swollen conidia, but no germlings in epithelial lung cells of leucopenic mice infected with *A. terreus* [6]. The precise role of alveolar epithelial cells in *A. terreus* infection remains unclear.

In this study, we used a micropatterned infection model recently developed in our laboratory [27] to investigate Arp2/3-dependent internalization of *A. terreus* conidia into phagolysosomes of alveolar epithelial cells. Our results provide novel insights into the molecular mechanisms underlying fungal internalization and persistence.

## Results

### Binding of *A. terreus* conidia to micropatterned cells

To investigate the interaction between *A. terreus* conidia and alveolar epithelial cells, we used a novel infection model based on micropatterned substrates, recently established and validated in our laboratory. Confining cells on micropatterns facilitates conidia internalization into phagosomes as compared to other cellular models, such as a large-scale cellular monolayer [27]. It also allows accurate mapping and quantification of conidia-cell interactions using an overlay of multiple identical micropatterns. Specifically, we cultured A549 epithelial cells that stably express phagolysosomal marker Lamp1-NeonGreen on fibronectin-coated circular micropatterns with a diameter of 85 µm. Under these conditions, cells consistently formed well-defined multicellular islands containing approximately 15–20 cells per micropattern (Fig 1A). Cells were further infected with dormant conidia of clinical isolate *A. terreus* sensu stricto for 1 and 3 hours. Infection resulted in conidia binding specifically to cellular islands, without unspecific adhesion observed on the surrounding substrate, confirming the high specificity and reproducibility of this infection model.

**Figure 1.**
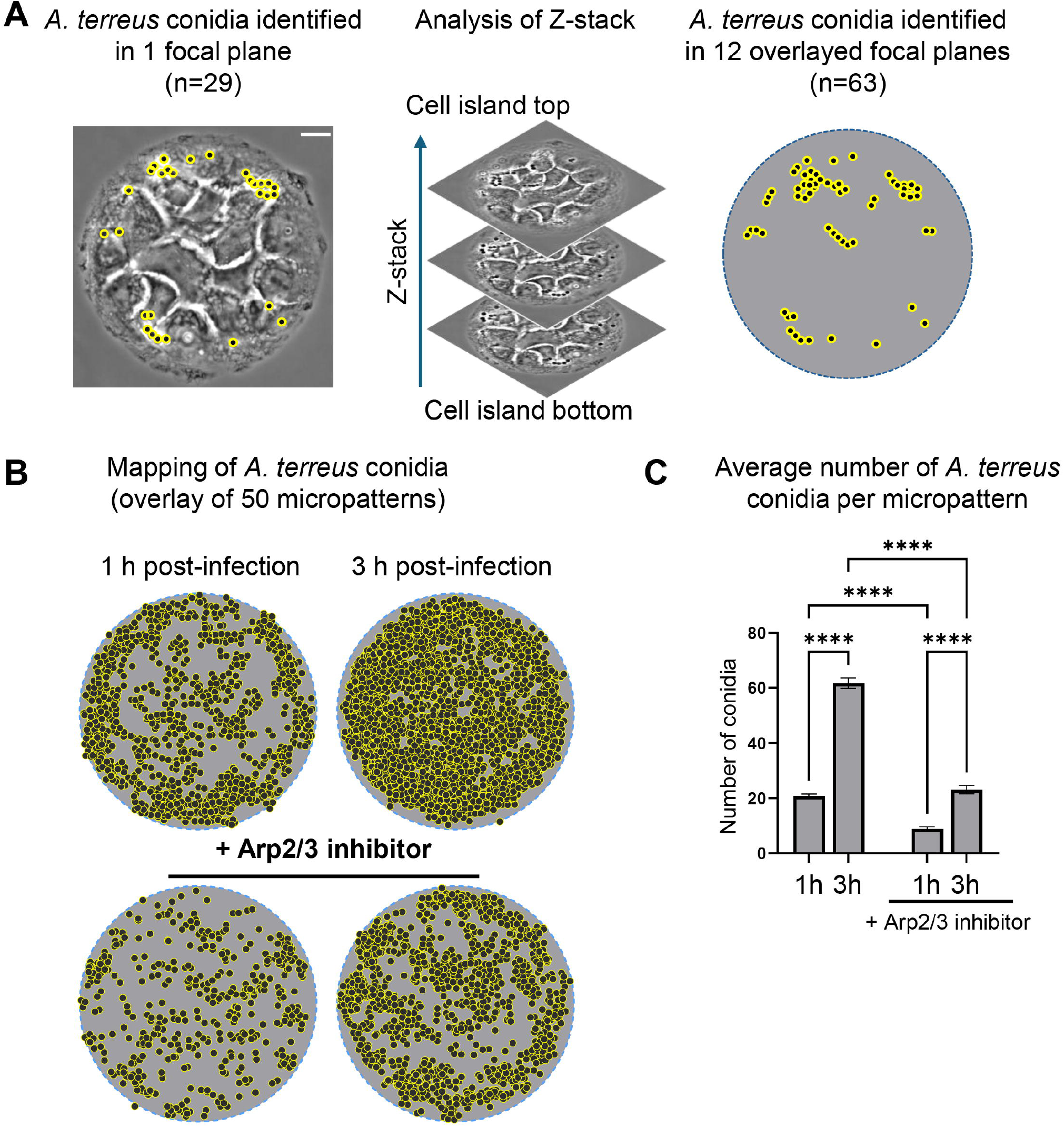
Arp2/3-dependent adhesion of *A. terreus* conidia to micropatterned alveolar epithelial cells. (A) Phase contrast image of A549 cells grown on a circular fibronectin-coated micropattern (85 µm diameter), forming a multicellular island. Shown is conidial identification (black dots with yellow circles) in a single focal plane versus 12 overlaid planes of Z-stack after infection with *A. terreus* conidia. Scale bar: 10 µm. (B) Computational mapping of conidial positions within 50 overlayed micropatterns. Each dot represents an individual detected conidium across the full Z-stack. (C) Quantification of average conidial number per micropattern under control and CK-666 conditions at 1 hour and 3 hours post-infection. Data represent mean ± SEM. ****p ≤ 0.0001.

To provide a quantitative basis for further analysis, we first quantified the total number of conidia attached to micropatterned cells at both infection time points. This assessment was conducted using multiplane phase-contrast microscopy, enabling quantification of conidia in 12 focal planes of a Z-stack (Fig 1A). More conidia could be identified when analysing a Z-stack as compared to a single focal plane. Phase contrast visualization included both surface-bound and internalized conidia, without discriminating between these states. Establishing the total conidia count per micropattern served further as an essential baseline metric. The subsequent internalization data were normalized to this number, ensuring accurate comparative analyses. Quantification demonstrated conidia binding to micropatterned cells already at 1 hour post-infection, with a significant further increase in conidia number observed after 3 hours (Fig 1B and Fig 1C, Table in S1 Table). Pre-treatment of cells with Arp2/3 inhibitor CK-666 significantly reduced the average number of total conidia associated with micropatterned cells at both the 1-hour and 3-hour time points (Fig 1B and Fig 1C).

### Co-localization of *A. terreus* conidia with actin-rich vesicles

To investigate the role of actin in fungal infection, we applied actin staining and quantified conidia association with actin-rich vesicular structures (Fig 2A). Given the inherent variability in conidia numbers bound per individual micropattern, normalization to the total conidia count per micropattern was performed to enable accurate comparative analysis between samples. Our quantitative evaluation revealed that conidial association with actin-rich vesicles (Actin^+^ vesicles) occurred rapidly and efficiently (Fig 2C and Fig 2D). Already after 1 hour of infection, 19.3% of *A. terreus* conidia were clearly associated with Actin^+^ vesicles. Interestingly, this percentage remained stable and did not significantly increase after 3 hours of infection. Notably, inhibition of the Arp2/3 complex dramatically decreased the number and percentage of conidia associated with Actin^+^ vesicles at 1 hour (1.4%), with some increase detected by 3 hours post-infection (7.6%) (Fig 2C and Fig 2D, Table in S1 Table).

**Figure 2.**
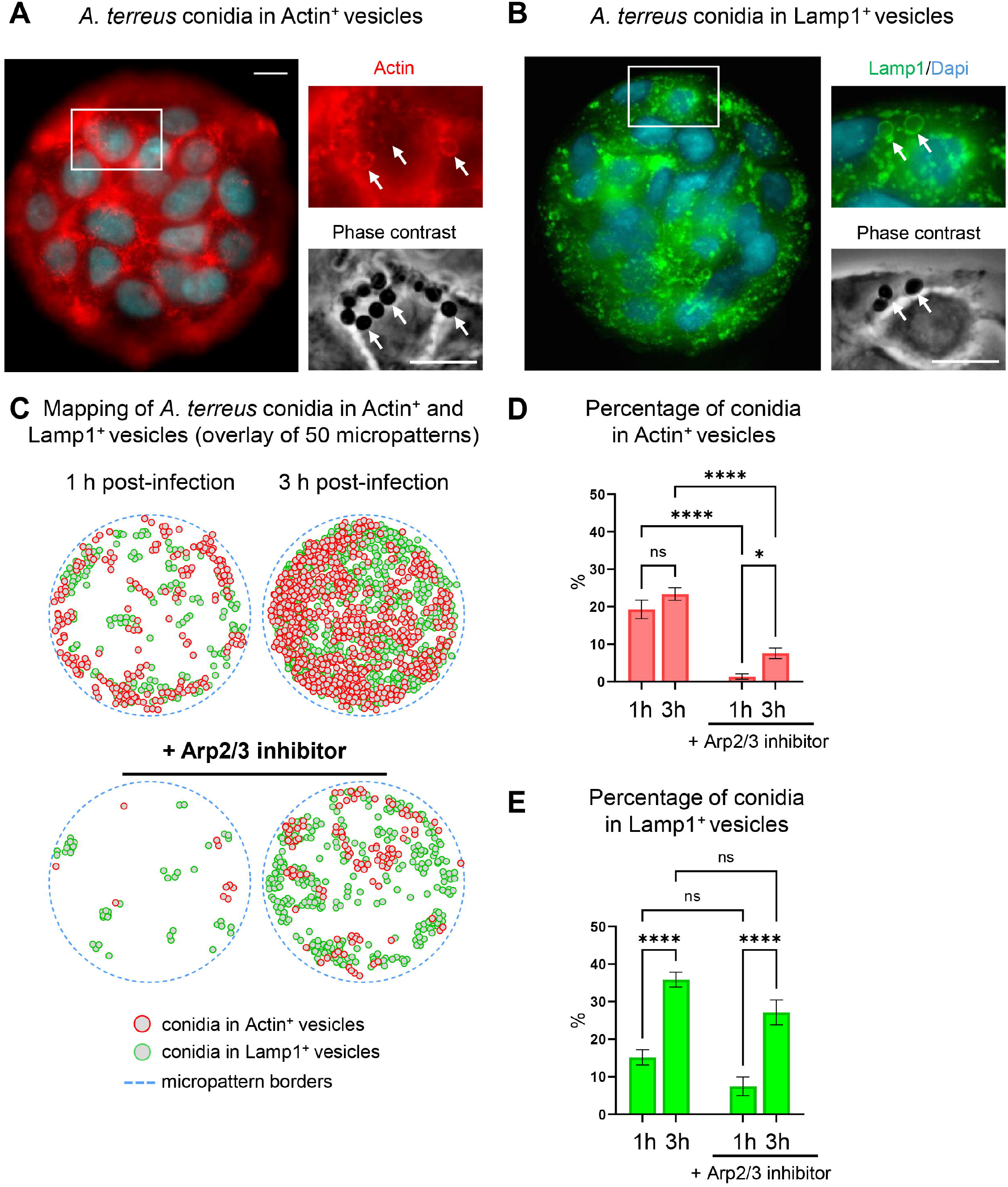
Internalization of *A. terreus* conidia into micropatterned alveolar epithelial cells. (A) Fluorescence image of micropatterned A549 cells stained with LifeAct to visualize actin (red), DAPI for nuclei (blue), and phase contrast image (grey). White arrows indicate conidia co-localized with actin structures in both TRITC and phase contrast channels. (B) Fluorescence image of micropatterned A549 cells expressing Lamp1-NeonGreen. Lamp1 (green), DAPI (blue), and phase contrast (grey). White arrows point to conidia localized within Lamp1^+^ vesicles. (C) Computational mapping of Actin^+^ (red circles) and Lamp1^+^ (green circles) vesicle positions within 50 overlayed micropatterns. Each circle represents an individual detected vesicle across the full Z-stack. (D) Quantification of the percentage of conidia in Actin^+^ vesicles. (E) Quantification of the percentage of conidia in Lamp1^+^ vesicles. Data represent mean ± SEM. *p ≤ 0.05; ****p ≤ 0.0001; ns, not significant; h-hour. Scale bar: 10 µm.

### Internalization of conidia into Lamp1-positive vesicles

To further dissect the intracellular fate of *A. terreus* conidia following epithelial internalization, we next examined their co-localization with Lamp1-positive phagolysosomes using Lamp1-NeonGreen as a marker (Fig 2B) [27,28]. Similarly to Actin^+^ vesicles, Lamp1 co-localization was quantified relative to the total number of conidia present on individual micropatterns. We observed that conidia internalization into Lamp1-positive phagolysosomes (Lamp1^+^ vesicles) occurred already after 1 hour of infection (Fig 2C and Fig 2E, Table in S1 Table). The percentage of conidia within Lamp1^+^ vesicles significantly increased between 1 hour (15.2%) and 3 hours (35.9%) (Fig 2E). Treatment of cells with Arp2/3 inhibitor substantially decreased the total number of conidia found in Lamp1^+^ vesicles (Fig 2C lower panel). However, when normalized relative to the total conidia count per micropattern, no statistically significant differences in the proportion of conidia in Lamp1^+^ vesicles were observed upon Arp2/3 inhibition (Fig 2E).

### Co-localization of actin patches with Lamp1-positive vesicles

While analysing the localization of *A. terreus* conidia within Lamp1^+^ phagolysosomes, we observed distinct actin accumulations on a subset of Lamp1^+^ vesicles containing conidia (Actin^+^Lamp1^+^ vesicles) (Fig 3A, Table in S1 Table). Actin patches that might resemble actin “flashes”, localized on top of Lamp1^+^ vesicles rather than co-localized with it. Actin^+^Lamp1^+^ vesicles were detected in 25 out of 50 analysed micropatterns already after 1 hour of infection (8.96%) and showed no significant change at 3 hours (7.64%) (Fig 3B, Fig 3C and Fig 3D, Table in S1 Table). Treatment with Arp2/3 inhibitor completely abolished actin patches at 1 hour, with only a minimal reappearance at 3 hours post-infection (2.17%) (Fig 3B and Fig 3C). Further, we calculated the fraction (ratio) of Actin^+^Lamp1^+^ vesicles relative to all identified Lamp1^+^ vesicles, revealing a decrease from 0.49 at 1 hour to 0.22 at 3 hours post-infection (Fig 3D). Following inhibitor treatment, the ratio quantification was only possible at the 3-hour time point and was not significant due to a very low abundance of Actin^+^Lamp1^+^ vesicles.

**Figure 3.**
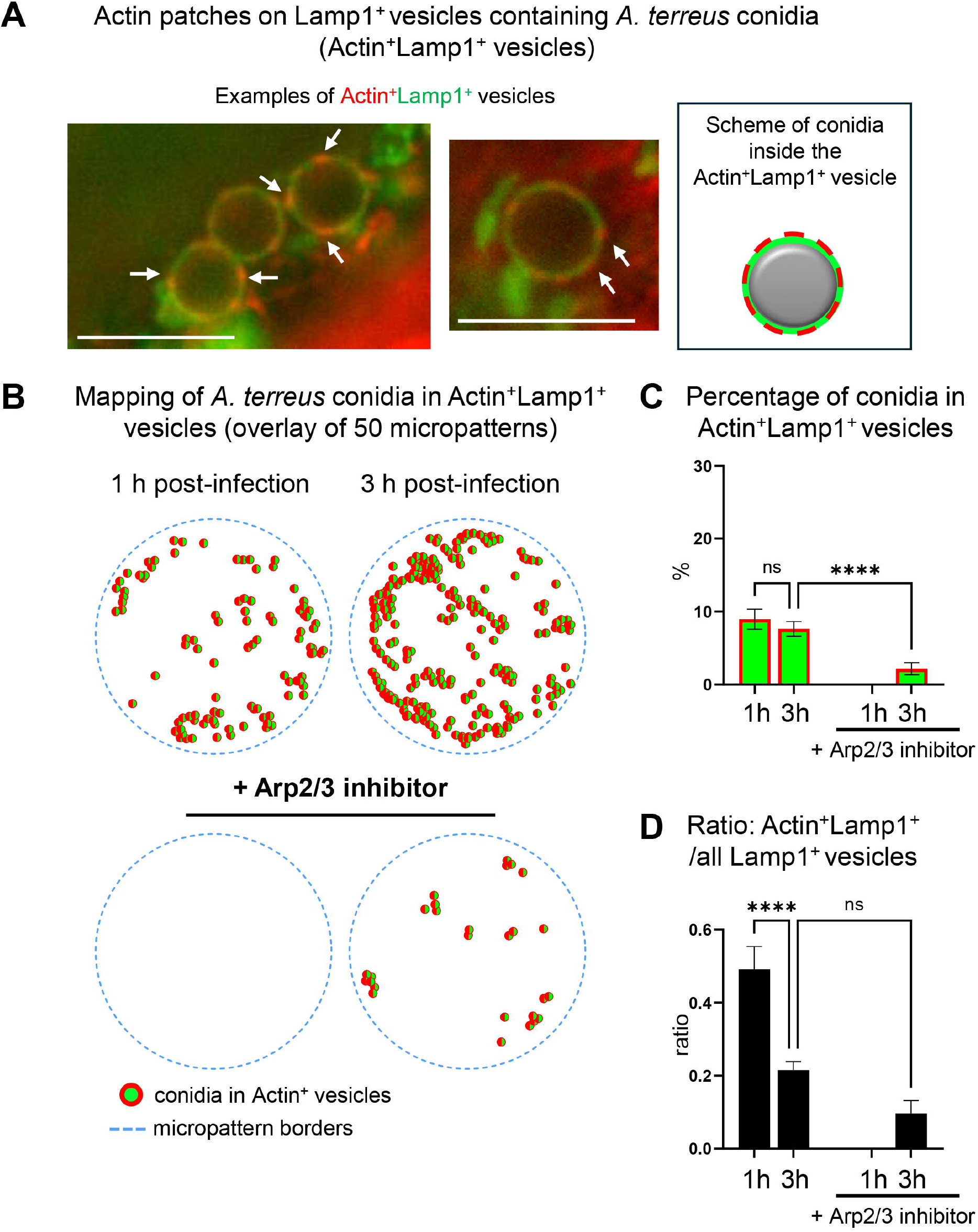
Actin patches decorate a subpopulation of Lamp1^+^ vesicles during *A. terreus* infection. (A) Representative fluorescence image showing Lamp1^+^ vesicles with localized actin patches (Actin^+^Lamp1^+^ vesicles). Actin (red), Lamp1 (green). White arrows indicate distinct actin accumulations. Scale bar: 5 µm. (B) Computational mapping of Actin^+^Lamp1^+^ vesicle positions within 50 overlayed micropatterns. Each circle represents an individual detected vesicle across the full Z-stack. (C) Quantification of conidia associated with Actin^+^Lamp1^+^ vesicles. (D) Ratio of double-positive vesicles (Actin^+^Lamp1^+^) relative to all identified Lamp1^+^ vesicles. Data represent mean ± SEM. ****p ≤ 0.0001; ns, not significant; h-hour.

## Discussion

In the present study, we investigated the early interactions of *A. terreus* conidia with alveolar epithelial cells. The primary objective was to elucidate processes of fungal adhesion and subsequent phagolysosomal internalization. These aspects were previously unexplored for *A. terreus* in epithelial models. Our findings confirm that epithelial internalization of *A. terreus* can be mechanistically dissected and offer new opportunities to explore host-pathogen interactions in detail.

Our findings revealed rapid adhesion of *A. terreus* conidia to micropatterned alveolar epithelial cell islands, detectable already 1 hour post-infection. The number of adherent conidia increased further by three hours, highlighting ongoing adhesion events. We also identified a central role of the Arp2/3 complex in mediating early interactions between epithelial cells and *A. terreus* conidia. Conidial adhesion was strongly dependent on Arp2/3-driven actin polymerization, consistent with its well-characterized function in supporting phagocytosis [26,29–32]. Inhibition of Arp2/3 led to a pronounced decrease in fungal binding and formation of Actin^+^ vesicles, indicating that actin polymerization is essential for establishing early host-pathogen contacts and initiating internalization. These observations also suggest that Arp2/3 may function as a dual mediator by facilitating both membrane-cytoskeleton coupling during uptake and cytoplasmic remodeling events post-entry.

Our study highlights the central importance of the actin cytoskeleton in the internalization of *A. terreus* conidia by alveolar epithelial cells. The rapid association of fungal conidia with actin-rich vesicles underlines actin polymerization as a crucial step in early internalization events. In professional phagocytes, Arp2/3-driven actin filament branching was shown to promote membrane protrusions and phagocytic cup formation, necessary for pathogen engulfment [31,33–35]. Our results extend previous findings to *A. terreus* and reveal that alveolar epithelial cells utilize Arp2/3-dependent mechanisms for fungal uptake.

A particularly interesting feature of the intracellular trafficking of *A. terreus* conidia was the differential behavior of Actin^+^ versus Lamp1^+^ vesicle populations over time. The Lamp1^+^ vesicles containing *A. terreus* conidia progressively accumulated between one and three hours of infection, supporting the ongoing phagosome maturation process. However, the percentage of conidia surrounded by actin remained comparatively stable. These results suggest that actin association is not a cumulative process, but rather a transient interaction that resolves quickly after pathogen internalization. We hypothesize that actin polymerization around internalized conidia is an early and short-lived event, potentially functioning as a dynamic scaffold to stabilize conidia binding. We have also observed a subset of Lamp1^+^ vesicles surrounded by distinct patches of actin, forming a population of Actin^+^Lamp1^+^ vesicles. These were not independent vesicles but rather Lamp1^+^ compartments transiently decorated with actin foci, morphologically consistent with actin flashes [36,37]. Actin flashes have been previously reported in phagocytic systems as short-lived, Arp2/3-mediated events that transiently associate with maturing phagosomes [38,36,37]. The function of actin flashes is not completely understood; they have been shown to modulate vesicle trafficking by controlling the timing of fusion events and/or limiting premature acidification [39,36,40]. In our system, the presence of actin-decorated Lamp1^+^ vesicles at both one and three hours of infection, as well as their strong sensitivity to Arp2/3 inhibition, supports the view that actin flashes may serve as regulatory checkpoints during phagosome maturation in epithelial cells. Moreover, the dissociation between overall conidial internalization and the frequency of Actin^+^Lamp1^+^ vesicles further supports this interpretation. As the total number of internalized conidia increased over time, Lamp1^+^ vesicle abundance rose accordingly. In contrast, the percentage of Actin^+^Lamp1^+^ vesicles did not increase, suggesting that actin recruitment to Lamp1^+^ phagosomes is temporally restricted and not a default outcome of maturation. Instead, actin flashes may occur on a subset of phagosomes. Their formation requires active Arp2/3 signaling, since we observed nearly complete loss upon inhibitor treatment.

Interestingly, a recent study has shown that the early endosomal GTPase Rab5c protein is critical for efficient phagosome maturation and LC3-associated phagocytosis in response to *A. fumigatus* conidia but is not required for their initial internalization [41]. This supports the view that uptake and intracellular processing are mechanistically separable. In our study, *A. terreus* conidia remained associated with Actin^+^ vesicles over time, suggesting delayed vesicle maturation. We can speculate that prolonged actin retention may delay the Rab5-dependent recruitment of early endosomal proteins and autophagy machinery, such as LC3, PI(3)P, and V-ATPase. This, in turn, could delay phagolysosomal fusion and acidification, contributing to the persistence of viable conidia in epithelial cells. Whether *A. terreus* actively modulates Rab5 signaling remains to be evaluated.

These findings introduce a new factor to the host response to *A. terreus* conidia. In contrast to *A. fumigatus*, which shows minimal actin co-localization under similar conditions [27] and our unpublished data), *A. terreus* induces a stronger actin response. The presence of Lamp1^+^ vesicles with actin flashes could reflect a fungal strategy to weaken normal degradation pathways, either by delaying phagosome maturation or by modulating antigen presentation mechanisms. The observed accumulation of conidia in Lamp1^+^ vesicles over time in untreated cells is consistent with progressive phagosome maturation toward degradative endpoints [27,22,42]. However, the ability of conidia to reach Lamp1^+^ vesicles even under Arp2/3 inhibition suggests that phagolysosomal routing is at least partially Arp2/3-independent, as described for other microbial pathogens [43,44]. Our results imply that once internalized, conidia can engage alternative trafficking routes.

In conclusion, our study provides the first mechanistic insights into *A. terreus* internalization and intracellular trafficking in alveolar epithelial cells. Our findings suggest an important role for the Arp2/3 complex in cell-conidia interactions. The formation of transitional compartments points to intracellular survival strategies of *A. terreus* to promote persistence. This might contribute to the clinical observation of delayed disease presentation. Further studies are needed to understand the molecular pathways underlying conidial internalization and to provide a basis for developing specific antifungal therapies.

## Materials and methods

### Fungal strain and growth conditions

The clinical isolate *Aspergillus terreus* sensu stricto (strain no 375) was cultured on Sabouraud 4% glucose agar (Sigma-Aldrich, 84088, Austria), prepared with 40 g/L D (+)-glucose (Sigma-Aldrich, 1.08337, Austria) and 10 g/L peptone (Carl Roth, 8986.1, Germany) and incubated at 37°C for ten days to ensure conidia maturation. Conidia were harvested by gently flooding the fungal culture surface with sterile spore buffer containing 0.1% Tween (Fisher Scientific, 10113103, Germany) and 0.9% NaCl (Sigma-Aldrich, 1.06404, Austria). The suspension was centrifuged at 4000 rpm for 5 min, the pellet was resuspended in sterile spore buffer, and was directly used for infection experiments.

### Cell culture and Arp2/3 inhibition

A549 human alveolar epithelial-like cell line stably expressing the phagolysosomal marker Lamp1-NeonGreen [28] was cultured in RPMI-1640 (Capricorn, RPMI-XA, Germany) supplemented with 10% fetal bovine serum (FBS, Capricorn Scientific, FBS-12A, Germany), 2 µl/ml puromycin, (Carl Roth, 0240.2, Germany), 1% penicillin/streptomycin mix (Capricorn Scientific, PS-B, Germany) and 1% L-glutamine (Capricorn Scientific, GLN-B, Germany). For Arp2/3 inhibition, cells were pretreated with the Arp2/3 complex inhibitor CK666 (150 µM, THP Medical Products, HY-16926, Austria) two hours prior to infection. The inhibitor was maintained throughout the experiment to ensure continuous disruption of Arp2/3-mediated actin polymerization, as established in previous studies [26].

### Micropatterned substrate preparation and cell seeding

Micropatterned substrates were fabricated on PEG-silane-modified glass coverslips using deep ultraviolet (UV) photolithography, as previously described [27,29,30]. Briefly, glass coverslips were incubated at 80 °C under an inert atmosphere in a 1 mM solution of linear PEG-silane (Rapp Polymere, Germany) dissolved in dry toluene for 20 hours, promoting covalent attachment of PEG to glass. PEG-coated coverslips were thoroughly rinsed with isopropanol, methanol, and distilled water, and dried with nitrogen. PEG-coated coverslips were placed on a chromium photomask featuring circular shapes of 85 µm in diameter (Compugraphics, Jena) and exposed to deep UV for 5 minutes (low-pressure mercury UV lamp, 185 nm, Heraeus Noblelight, Germany) at 5 cm distance. The UV exposure locally oxidized the PEG layer, allowing subsequent protein adsorption. Micropatterned coverslips were further incubated at 4 °C overnight with fibronectin (10 µg/ml in PBS, Sigma-Aldrich, 341631, Austria) to promote cell adhesion. After rinsing in PBS, cells were seeded at a density of 1×10^6^ cells/mL in RPMI with 1% FBS and incubated for 2 hours at 37 °C. Non-adherent cells were gently removed by washing with pre-warmed PBS. The remaining cells were cultured overnight in the same serum-reduced (1%) RPMI medium to allow proper spreading on micropatterns.

### Infection with *A. terreus* conidia

After overnight incubation on micropatterned fibronectin-coated coverslips, A549 cells were infected with 10^7^ cfu/ml of *A. terreus* conidia. Before infection, serum-reduced medium was replaced with full-growth RPMI-1640 containing 10% FBS and no antibiotics. Cells were incubated with conidia for either 1 hour or 3 hours at 37 °C under standard cell culture conditions and further processed for fixation.

### Fluorescence microscopy

To preserve Lamp1-positive vesicles and actin cytoskeleton, cells were fixed using a “mild fixation” protocol, specifically optimized for these structures [31]. Fixed cells were further incubated with 0.05% saponin (Sigma-Aldrich, 47036, Austria) in PBS for permeabilization and simultaneously stained with EasyProbe ™ ActinRed 555 (FP032, THP Medical Products, Austria) and DAPI (1:20.000, D9542-5MG, Sigma-Aldrich, Austria) at room temperature for 1 hour. Following three PBS washes, samples were mounted on glass slides using Mowiol mounting medium (Sigma-Aldrich C9368, Austria). Fluorescence images were acquired using an Olympus IX83 inverted wide-field microscope (Olympus Austria) equipped with a UPlanXApo 60×/1.42 NA objective and 2X digital zoom. Z-stacks were recorded in parallel using a 0.5 µm step size across all imaging channels (DAPI, FITC, TRITC, and phase contrast PH3), at a resolution of 2048×2048 pixels and 16-bit grayscale. Exposure times were 8 ms (DAPI), 120–150 ms (TRITC/FITC), and 60 ms (phase contrast). Image stacks were deconvoluted using Olympus cellSens software with nearest-neighbor settings (50%). For each condition, 50 micropatterned cells from three biological replicates were analysed.

### Image analysis and quantification

Image evaluation was carried out using Fiji software (RRID: SCR_002285). Three trained researchers independently identified and quantified vesicle types by analysing merged fluorescence channels: Actin^+^ (TRITC), Lamp1^+^ (FITC), and Lamp1^+^Actin^+^ (TRITC/FITC co-localization) compartments. The “Point tool” and “Analyze-Measure” functions were used to record the X/Y-positions of fungal conidia and their associated vesicles. Z-stack data were reviewed in parallel to assess cell numbers (nuclear staining with DAPI), total conidial counts (phase contrast), and subcellular localization in relation to actin and Lamp1 fluorescent signals.

### Statistical analysis

Quantitative data were processed in GraphPad Prism 10.1.2 (RRID: SCR_002798). Statistical tests included descriptive statistics and one-way ANOVA with multiple comparisons. Figures were prepared in Fiji and finalized using Adobe Photoshop (RRID: SCR_014199). The statistical significance of the data was verified by p-values.

## Supporting information

**S1 Table. Summary of quantitative analysis of A. terreus conidia interactions with micropatterned A549 cells.**

## References

1. WHO Antimicrobial Resistance Division (2022) WHO fungal priority pathogens list to guide research, development and public health action. Available: https://iris.who.int/bitstream/handle/10665/363682/9789240060241-eng.pdf?sequence=1. Accessed 11 September 2024.

2. Bongomin F, Gago S, Oladele RO, Denning DW (2017) Global and Multi-National Prevalence of Fungal Diseases-Estimate Precision. Journal of fungi (Basel, Switzerland) 3 (4).

3. Lass-Flörl C (2012) Aspergillus terreus: how inoculum size and host characteristics affect its virulence. The Journal of infectious diseases 205 (8): 1192–1194.

4. Lass-Flörl C (2018) Treatment of Infections Due to Aspergillus terreus Species Complex. Journal of fungi (Basel, Switzerland) 4 (3).

5. Perkhofer S, Lass-Flörl C, Hell M, Russ G, Krause R et al. (2010) The Nationwide Austrian Aspergillus Registry: a prospective data collection on epidemiology, therapy and outcome of invasive mould infections in immunocompromised and/or immunosuppressed patients. International journal of antimicrobial agents 36 (6): 531– 536.

6. Slesiona S, Ibrahim-Granet O, Olias P, Brock M, Jacobsen ID (2012) Murine infection models for Aspergillus terreus pulmonary aspergillosis reveal long-term persistence of conidia and liver degeneration. The Journal of infectious diseases 205 (8): 1268–1277.

7. Lass-Flörl C, Rath P, Niederwieser D, Kofler G, Würzner R et al. (2000) Aspergillus terreus infections in haematological malignancies: molecular epidemiology suggests association with in-hospital plants. The Journal of hospital infection 46 (1): 31–35.

8. Lass-Flörl C, Lackner M (2018) Azole-Resistance in Aspergillus terreus and Related Species: An Emerging Problem or a Rare Phenomenon. Frontiers in microbiology (9): 516.

9. Lass-Flörl C, Dietl A-M, Dimitrios P. Kontoyiannis, Dimitrios P., Brock M (2021) Aspergillus terreus Species Complex. Clinical Microbiology Reviews 43 (4).

10. Vahedi-Shahandashti R, Lass-Flörl C (2019) Antifungal resistance in Aspergillus terreus: A current scenario. Fungal genetics and biology : FG & B 131: 103247.

11. Thakur R, Shishodia SK, Sharma A, Chauhan A, Kaur S et al. (2024) Accelerating the understanding of Aspergillus terreus: Epidemiology, physiology, immunology and advances. Current research in microbial sciences 6: 100220.

12. Lass-Flörl C, Griff K, Mayr A, Petzer A, Gastl G et al. (2005) Epidemiology and outcome of infections due to Aspergillus terreus: 10-year single centre experience. British journal of haematology 131 (2): 201–207.

13. Ibrahim-Granet O, Philippe B, Boleti H, Boisvieux-Ulrich E, Grenet D et al. (2003) Phagocytosis and intracellular fate of Aspergillus fumigatus conidia in alveolar macrophages. Infection and immunity 71 (2): 891–903.

14. Chignard M, Balloy V, Sallenave J-M, Si-Tahar M (2007) Role of Toll-like receptors in lung innate defense against invasive aspergillosis. Distinct impact in immunocompetent and immunocompromized hosts. Clinical immunology (Orlando, Fla.) 124 (3): 238–243.

15. Luther K, Torosantucci A, Brakhage AA, Heesemann J, Ebel F (2007) Phagocytosis of Aspergillus fumigatus conidia by murine macrophages involves recognition by the dectin-1 beta-glucan receptor and Toll-like receptor 2. Cellular microbiology 9 (2): 368– 381.

16. Slesiona S, Gressler M, Mihlan M, Zaehle C, Schaller M et al. (2012) Persistence versus escape: Aspergillus terreus and Aspergillus fumigatus employ different strategies during interactions with macrophages. PloS one 7 (2): e31223.

17. Chiang LY, Sheppard DC, Gravelat FN, Patterson TF, Filler SG (2008) Aspergillus fumigatus stimulates leukocyte adhesion molecules and cytokine production by endothelial cells in vitro and during invasive pulmonary disease. Infection and immunity 76 (8): 3429–3438.

18. Park SJ, Burdick MD, Brix WK, Stoler MH, Askew DS et al. (2010) Neutropenia enhances lung dendritic cell recruitment in response to Aspergillus via a cytokine-to-chemokine amplification loop. Journal of immunology (Baltimore, Md. : 1950) 185 (10): 6190–6197.

19. Heinekamp T, Thywißen A, Macheleidt J, Keller S, Valiante V et al. (2012) Aspergillus fumigatus melanins: interference with the host endocytosis pathway and impact on virulence. Frontiers in microbiology 3: 440.

20. Stappers MHT, Clark AE, Aimanianda V, Bidula S, Reid DM et al. (2018) Recognition of DHN-melanin by a C-type lectin receptor is required for immunity to Aspergillus. Nature 555 (7696): 382–386.

21. Hsieh S-H, Kurzai O, Brock M (2017) Persistence within dendritic cells marks an antifungal evasion and dissemination strategy of Aspergillus terreus. Scientific Reports 7 (1): 10590.

22. Wasylnka JA, Moore MM (2002) Uptake of Aspergillus fumigatus Conidia by phagocytic and nonphagocytic cells in vitro: quantitation using strains expressing green fluorescent protein. Infection and immunity 70 (6): 3156–3163.

23. Wasylnka JA, Moore MM (2003) Aspergillus fumigatus conidia survive and germinate in acidic organelles of A549 epithelial cells. Journal of cell science 116 (Pt 8): 1579–1587.

24. Bigot J, Guillot L, Guitard J, Ruffin M, Corvol H et al. (2020) Bronchial Epithelial Cells on the Front Line to Fight Lung Infection-Causing Aspergillus fumigatus. Frontiers in Immunology 11: 1041.

25. Osherov N (2012) Interaction of the pathogenic mold Aspergillus fumigatus with lung epithelial cells. Frontiers in microbiology 3: 346.

26. Culibrk L, Croft CA, Toor A, Yang SJ, Singhera GK et al. (2019) Phagocytosis of Aspergillus fumigatus by Human Bronchial Epithelial Cells Is Mediated by the Arp2/3 Complex and WIPF2. Frontiers in cellular and infection microbiology 9: 16.

27. Schiefermeier-Mach N, Polleux J, Heinrich L, Lechner L, Vorona O et al. (2025) Biological boundary conditions regulate the internalization of Aspergillus fumigatus conidia by alveolar cells. Frontiers in cellular and infection microbiology 15: 1515779.

28. Schiefermeier-Mach N, Moresco V, Geley S, Heinrich L, Lechner L et al. (2021) Evaluation of Stable LifeAct-mRuby2- and LAMP1-NeonGreen Expressing A549 Cell Lines for Investigation of Aspergillus fumigatus Interaction with Pulmonary Cells. International journal of molecular sciences 22 (11).

29. Castellano F, Chavrier P, Caron E (2001) Actin dynamics during phagocytosis. Seminars in immunology 13 (6): 347–355.

30. May RC, Caron E, Hall A, Machesky LM (2000) Involvement of the Arp2/3 complex in phagocytosis mediated by FcgammaR or CR3. Nature cell biology 2 (4): 246–248.

31. Rotty JD, Wu C, Bear JE (2013) New insights into the regulation and cellular functions of the ARP2/3 complex. Nature reviews. Molecular cell biology 14 (1): 7–12.

32. Vorselen D, Barger SR, Wang Y, Cai W, Theriot JA et al. (2021) Phagocytic ‘teeth’ and myosin-II ‘jaw’ power target constriction during phagocytosis. eLife 10.

33. Flannagan RS, Jaumouillé V, Grinstein S (2012) The cell biology of phagocytosis. Annual review of pathology 7: 61–98.

34. Freeman SA, Grinstein S (2014) Phagocytosis: receptors, signal integration, and the cytoskeleton. Immunological reviews 262 (1): 193–215.

35. Jaumouillé V, Waterman CM (2020) Physical Constraints and Forces Involved in Phagocytosis. Frontiers in Immunology 11.

36. Liebl D, Griffiths G (2009) Transient assembly of F-actin by phagosomes delays phagosome fusion with lysosomes in cargo-overloaded macrophages. Journal of cell science 122 (Pt 16): 2935–2945.

37. Poirier MB, Fiorino C, Rajasekar TK, Harrison RE (2020) F-actin flashes on phagosomes mechanically deform contents for efficient digestion in macrophages. Journal of cell science 133 (12).

38. Yam PT, Theriot JA (2004) Repeated cycles of rapid actin assembly and disassembly on epithelial cell phagosomes. Molecular biology of the cell 15 (12): 5647–5658.

39. Johnston SA, May RC (2010) The human fungal pathogen Cryptococcus neoformans escapes macrophages by a phagosome emptying mechanism that is inhibited by Arp2/3 complex-mediated actin polymerisation. PLoS Pathogens 6 (8): e1001041.

40. Patel DM, Ahmad SF, Weiss DG, Gerke V, Kuznetsov SA (2011) Annexin A1 is a new functional linker between actin filaments and phagosomes during phagocytosis. Journal of cell science 124 (Pt 4): 578–588.

41. Freitas-Filho EG, Zaidan I, Fortes-Rocha M, Alzamora-Terrel DL, Bifano C et al. (2025) RAB5c controls the assembly of non-canonical autophagy machinery to promote phagosome maturation and microbicidal function of macrophages. bioRxiv : the preprint server for biology.

42. Jia L-J, Rafiq M, Radosa L, Hortschansky P, Cunha C et al. (2023) Aspergillus fumigatus hijacks human p11 to redirect fungal-containing phagosomes to non-degradative pathway. Cell host & microbe 31 (3): 373-388.e10.

43. Epp Elias, Nazarova Elena, Regan Hannah, Douglas Lois M., Konopka James B. et al. (2013) Clathrin- and Arp2/3-Independent Endocytosis in the Fungal Pathogen Candida albicans. mBio 4 (5): 10.1128/mbio.00476-13.

44. Rotty JD, Brighton HE, Craig SL, Asokan SB, Cheng N et al. (2017) Arp2/3 Complex Is Required for Macrophage Integrin Functions but Is Dispensable for FcR Phagocytosis and In Vivo Motility. Developmental cell 42 (5): 498-513.e6. Available: https://www.sciencedirect.com/science/article/pii/S1534580717306317.

