## Supplemental Table 1 for "Arp2/3 complex contributes to the actin-dependent uptake of *Aspergillus terreus* conidia by alveolar epithelial cells"

### Supporting information

#### S1 Table. Summary of quantitative analysis of *A. terreus* conidia interactions with micropatterned A549 cells

| Time post-infection | Conidia in Actin$^+$ vesicles | Conidia in Lamp1$^+$ vesicles | Actin patches on Lamp1$^+$ vesicles (Actin$^+$Lamp1$^+$ vesicles) | Ratio of Actin$^+$Lamp1$^+$ to all Lamp1 vesicles | Overage number of conidia per micropattern | Total number of conidia (50 micropatterns) |
|---|---|---|---|---|---|---|
| 1 hour | 19.30±2.48% | 6.20±1.30% | 8.96±1.38% | 0.49±0.06 | 20.74±0.80 | 1037 |
| 3 hours | 23.40±1.67% | 28.20±1.76% | 7.64±1.00% | 0.22±0.02 | 61.72±1.90 | 3086 |
| 1hour (+CK666) | 1.35±0.69% | 7.47±2.52% | 0.00±0.00% | 0.00 | 8.84±0.76 | 442 |
| 3 hours (+CK666) | 7.59±1.40% | 25.0±3.29% | 2.17±0.832% | 0.10±0.04 | 23.14±1.52 | 1157 |
